# Biphasic impact of prenatal inflammation and macrophage depletion on the wiring of neocortical inhibitory circuits

**DOI:** 10.1101/669002

**Authors:** Morgane Sonia Thion, Coralie-Anne Mosser, Isabelle Férézou, Pauline Grisel, Sofia Baptista, Donovan Low, Florent Ginhoux, Sonia Garel, Etienne Audinat

## Abstract

The etiology of neurodevelopmental disorders is linked to defects in Parvalbumin (PV)-expressing cortical interneurons and to prenatal immune challenges. Mouse models of maternal immune activation (MIA) and microglia deficits increase the postnatal density of PV interneurons, raising the question of their functional integration. Here, we show that MIA and embryonic depletion of macrophages including microglia, have a two-step impact on PV interneurons wiring onto their excitatory target neurons in the barrel cortex. In adults, both challenges reduced the inhibitory drive from PV interneurons, as reported in neurodevelopmental disorders. In juveniles, however, we found an increased density of PV neurons, an enhanced strength of unitary connections onto excitatory cells and an aberrant horizontal inhibition with a reduced lateral propagation of sensory inputs *in vivo*. Our results provide a novel framework for understanding the impact of prenatal immune challenges onto the developmental trajectory of inhibitory circuits that leads to pathological brain wiring.

## INTRODUCTION

Prenatal inflammation and dysfunction of microglia, the brain resident macrophages, have both been associated with the etiology of several neuropsychiatric disorders, including schizophrenia (SZ) and autism spectrum disorders (ASD) (Beumer et al., 2012; Choi et al., 2016; Estes and McAllister, 2016; Hammond et al., 2018; Hanamsagar and Bilbo, 2017; Kim et al., 2017; Mattei et al., 2017; Patterson, 2011; Sekar et al., 2016; Shin Yim et al., 2017; Solek et al., 2018; Takano, 2015). However, how these prenatal immune challenges impact on the developmental trajectory of neocortical circuits remains to be fully examined. On the one hand, microglia invade the brain during early embryogenesis (Hoeffel and Ginhoux, 2018; Prinz et al., 2017; Thion et al., 2018) and constitute key actors of excitatory synapse density and function during postnatal and adult life (Li and Barres, 2018; Miyamoto et al., 2016; Paolicelli and Ferretti, 2017; Salter and Stevens, 2017; Schafer and Stevens, 2015; Wolf et al., 2017; Wu et al., 2015). On the other hand, models of prenatal inflammation using maternal immune activation (MIA) in rodents have been shown to induce in the adult offspring reduced functional GABAergic transmission from a specific population of interneurons expressing Parvalbumin (PV) in both the prefrontal and the somatosensory cortices (Canetta et al., 2016; Shin Yim et al., 2017). Inhibitory fast spiking (FS) PV interneurons act as gain modulators in neocortical circuits, play important roles in the plasticity and functional maturation of the neocortical network during development (Butt et al., 2017; Hensch, 2005). For instance, in sensory cortices, PV interneurons control feed-forward inhibition (FFI) in response to external stimuli, which shapes the temporal response in principal neurons, and is essential for sensory perception (Chittajallu and Isaac, 2010; Feldmeyer et al., 2018). In agreement with their important roles in cortical circuits, increasing evidence points towards PV interneuron dysfunction in neurodevelopmental psychiatric disorders (Nelson and Valakh, 2015). Remarkably, manipulating the activity of PV neurons in adults in the somatosensory or prefrontal cortex is sufficient to rescue some behavioral deficits triggered by MIA (Canetta et al., 2016; Shin Yim et al., 2017). We previously found that both MIA and microglia embryonic dysfunction modulate the early positioning of interneurons in the developing cortical plate with a subsequent mild increase in the density of PV interneurons in the murine somatosensory cortex (Squarzoni et al., 2014). These findings raise the possibility that prenatal immune challenges, including microglia deficits or MIA, which modifies microglial functioning but also a wide range of additional processes (Estes and McAllister, 2016; Smolders et al., 2018), might interfere with the developmental trajectory of PV interneurons from early stages.

Here, by combining anatomical, electrophysiological and imaging approaches, we show that transient early macrophage depletion and MIA lead to a reduced inhibitory drive of PV interneurons onto their targets in the layer 4 of the postnatal barrel cortex, consistent with what has been reported in other MIA paradigms. However, this hypo-inhibition was preceded by an unexpected, profound miswiring and hyperconnectivity of PV interneurons in juveniles, at a timepoint critical for learning and plasticity. These findings reveal a convergence of distinct immune challenges onto well-established actors in cortical networks and highlight a role of microglia in the wiring of inhibitory circuits. Importantly, they reveal an unexpected temporal trajectory, with major importance for understanding pathological brain wiring and designing effective interventional therapies.

## RESULTS

### Reduced inhibitory drive of PV interneurons onto principal cells in adult mice

To investigate how prenatal immune challenges impact on inhibitory circuits, we used genetically engineered mice to label PV interneurons (*PV*^*cre/+*^*;R26*^*tom/+*^) in two experimental paradigms: i) MIA triggered by a lipopolysaccharide (LPS) injection at embryonic day (E) 13.5, which mimics a bacterial infection; ii) early injections of anti-CSF-1R antibodies, which target macrophage precursors in the yolk-sac and lead to a severe depletion of microglia from E7 to E18.5, followed by a repopulation completed by postnatal day (P) 7 (Squarzoni et al., 2014; Waisman et al., 2015). Notably, our transient depletion model targets all embryonic macrophages, including microglia, meningeal macrophages, perivascular macrophages and other peripheral populations, which repopulate around E14.5 (Hoeffel et al., 2015). Thus, whereas brain-specific phenotypes are likely due to the depletion of microglia, a contribution of additional populations of macrophages cannot be excluded. We previously reported that both MIA and depletion models were shown to cause an increase in inhibitory interneurons entering the cortical plate (Squarzoni et al., 2014). Before probing the connectivity of PV interneurons, we carefully examined the overall barrel cortex architecture using the thalamocortical axonal marker, vGlut2 in coronal and tangential sections of flattened cortices (**Figure S1A-K**). After MIA or *in utero* macrophage depletion, we found a similar organization of the barrel field (**Figure S1A-K**) and an unchanged number of microglia at P7 compared to controls (**Figure S1L-M**), confirming that the microglial depletion is transient. We next examined both models at P60, in young adults, and found no significant change in the overall density of PV cells within the barrel cortex (**Figures 1A-B** and **S2A**). To examine functional properties, we recorded unitary inhibitory post-synaptic currents (uIPSC) evoked by PV FS interneurons onto principal excitatory neurons contained within the same barrel at P60-65 (**Figure 1C-D**). In both models, the amplitude of the unitary synaptic responses in principal cells (PC) was almost reduced by half compared to control mice (**Figure 1D-F**). The short-term dynamics of unitary IPSCs evoked by trains of presynaptic action potentials was not dramatically affected (**Figure 1F**), although a small but consistent decrease in paired-pulse ratio (PPR), consistent with a small increase in release probability, was observed (data not shown). These severe phenotypes on the amplitude of unitary IPCSs are consistent with previous findings reporting lower synaptic inhibition mediated by PV cells in the prefrontal or somatosensory cortices in the MIA offspring (Canetta et al., 2016; Shin Yim et al., 2017). Since microglia have been involved in synaptic formation and elimination (Miyamoto et al., 2016; Schafer and Stevens, 2015), we tested whether the sharp decrease in unitary synaptic responses could be explained by a major reduction in the number of synapses. We found a very limited or no decrease in the numbers of synaptic boutons formed by PV interneurons onto the soma of PC in the two experimental conditions (**Figure S3**). While this reduction could contribute to the observed electrophysiological phenotype, there are likely additional and complex changes in synaptic weight or composition in MIA and depleted offspring. Taken together, our experiments reveal a long-term impact of prenatal inflammation and microglia depletion onto the wiring of PV inhibitory circuits.

**Figure 1.**
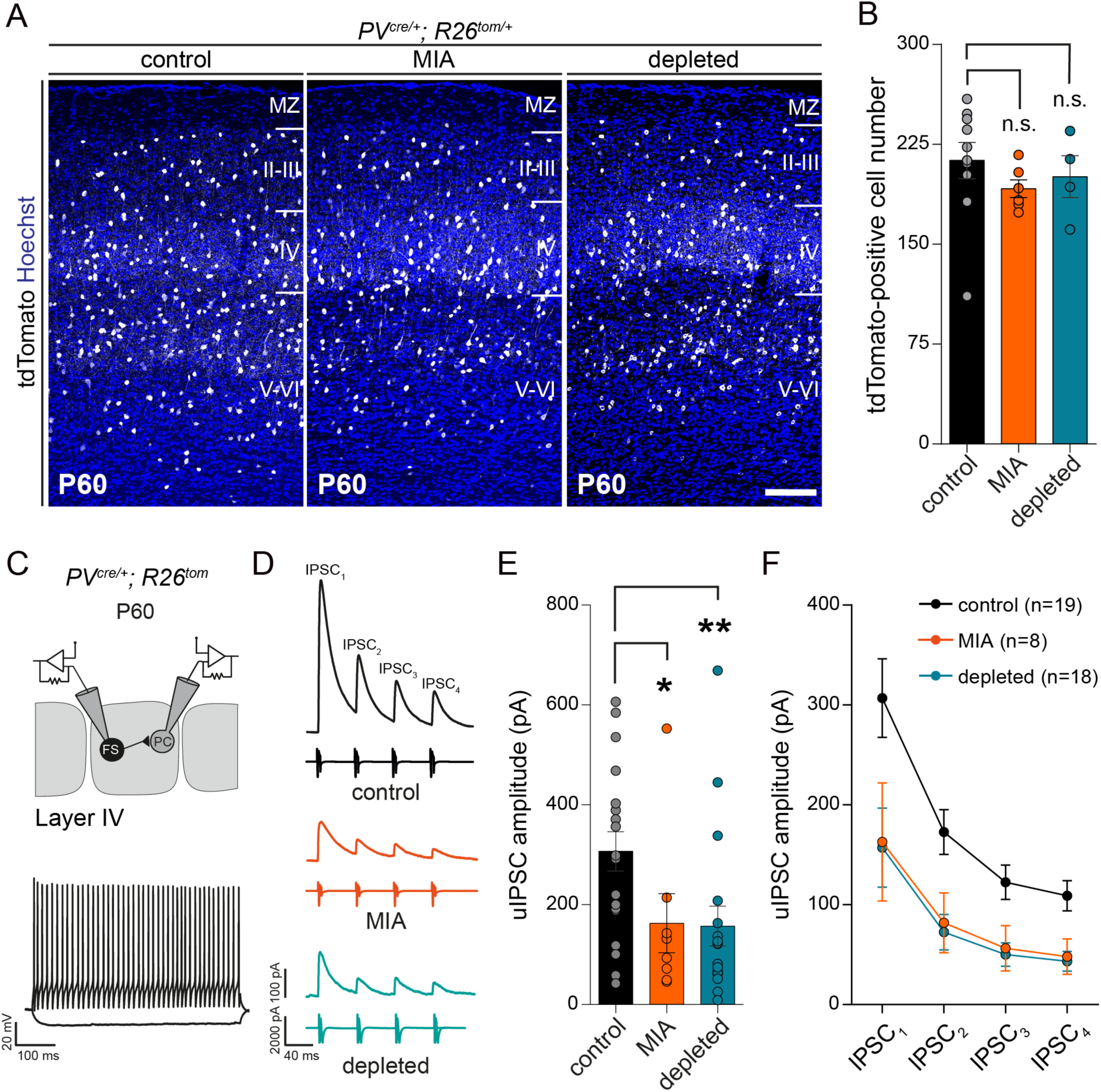
Maternal Immune Activation and *in utero* microglia depletion do not affect the density of Parvalbumin interneurons but decrease their synaptic connections onto principal neurons in layer 4 of the adult barrel cortex. **(A)** Coronal sections through the P60 somatosensory barrel cortex of control, MIA and microglia depleted *PV*^*Cre/+*^*;R26*^*tom/+*^ mice revealing no significant modulation of tdTomato-positive interneurons (control: N = 10; MIA: N = 6; depleted: N = 4). Scale bar equal = 150 µm. **(B)** Comparison of tdTomato-positive interneurons density within cortical layers between controls, MIA and depleted P60 mice (control: N = 10, MIA: N = 6, depleted: N = 4). **(C)** Scheme of the recording configuration depicting dual whole-cell recordings from a fast-spiking PV interneuron (FS) targeting a layer 4 principal cell (PC) in the barrel cortex. The inset shows representative voltage traces of a FS cell in response to hyperpolarizing and depolarizing current pulses. **(D)** Representative average traces of inhibitory postsynaptic currents (IPSC_1-4_) evoked in PC by trains of action potentials elicited in FS interneurons in thalamocortical slices of control, MIA and microglia-depleted P60 mice. **(E)** Comparison of the first IPSC (IPSC_1_) amplitude (control: N = 6, n_cells_ = 12; MIA: N = 4, n_cells_ = 8; depleted: N = 3, n_cells_ = 8). **(F)** Comparison of the IPSC amplitude evolution during the train between the three experimental conditions. Data are represented as mean ± SEM. Two-sided unpaired Mann-Whitney test was performed to assess differences. *p < 0.05, **p < 0.01, n.s., not significant. See also Figure S2.

### Enhanced inhibitory drive of PV interneurons onto principal cells in juveniles

To explore further the developmental trajectory of the PV network, we focused on P20, which corresponds to a timepoint when neurons express robust levels of PV and to a key period of learning an exploration. At this stage, we detected a mild increase in the density of PV neurons after MIA or transient embryonic depletion of microglia (**Figures 2A-B** and **S2A**). Next, we recorded unitary inhibitory synaptic connections made by PV FS interneurons onto PC in the same barrel at P20-25 (**Figure 2C-D**). Notably, the firing properties of these FS interneurons was not altered by experimental conditions (**Figure 2C** and data not shown). In sharp contrast to what we observed in adult mice, the amplitude of unitary synaptic responses was almost twice as large in both MIA and embryonically depleted pups compared to controls (**Figure 2C-F**). The short-term dynamics of unitary IPSCs evoked by trains of presynaptic action potentials was not dramatically affected (**Figure 2F**), although a small but consistent decrease in paired-pulse ratio, consistent with a small increase in release probability, was observed (data not shown). In addition, the densities of synaptic boutons formed by PV interneurons onto the soma of PC was only slightly increased (**Figure S3**). These observations thus indicate that the observed phenotype is likely due to complex modifications with several contributing factors (**Figure S3**). Irrespective of the underlying mechanisms, our results show that immune challenges during prenatal life induce, in juveniles, a transient hyper-inhibition of PC by PV interneurons in layer 4.

**Figure 2.**
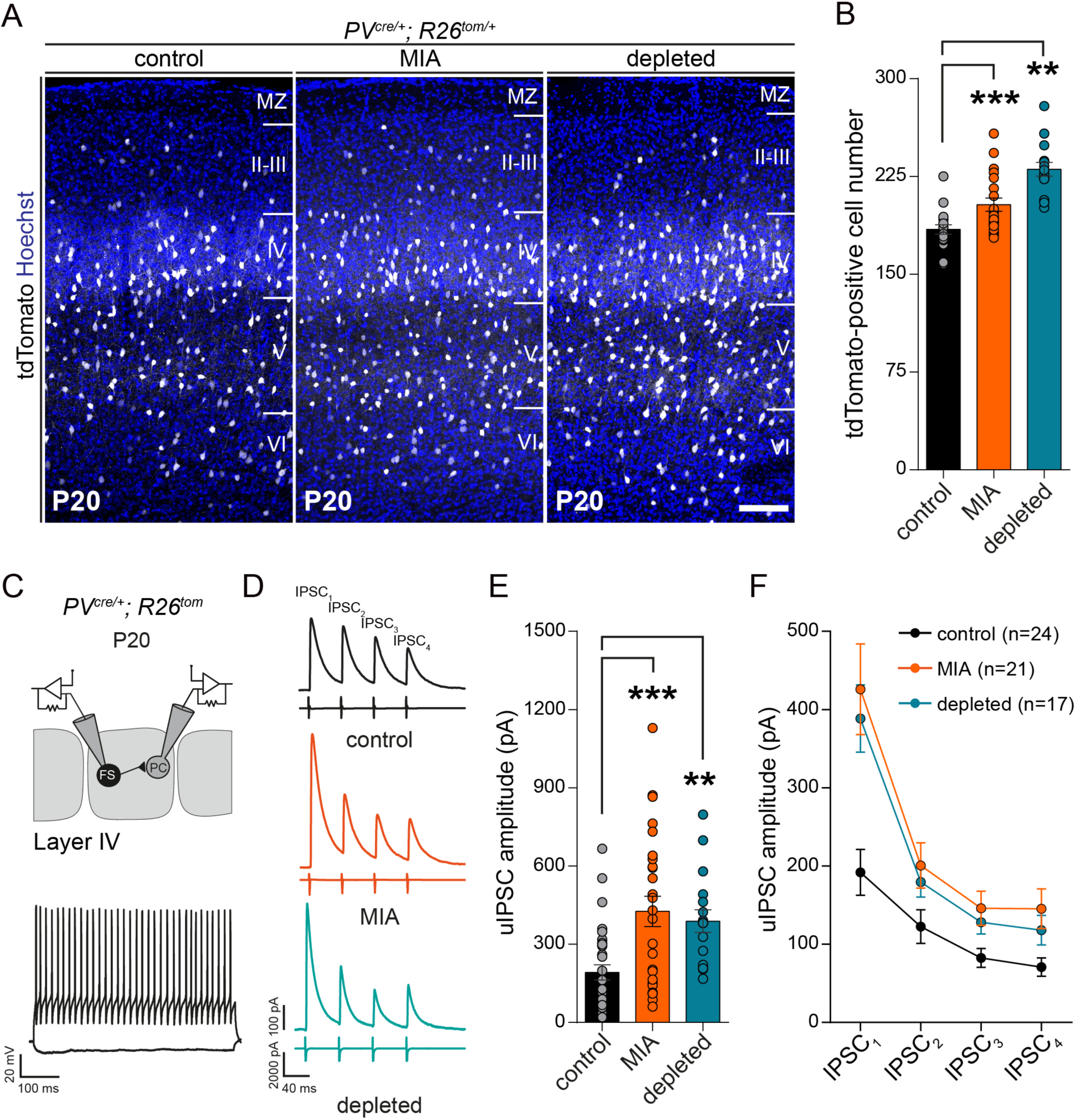
Maternal immune activation and *in utero* microglia depletion alter the density of PV interneurons and enhance their synaptic connections onto principal neurons in layer 4 of the juvenile barrel cortex. Coronal sections through the barrel cortex of control, MIA and microglia depleted P20 *PV*^*Cre/+*^*;R26*^*tom/+*^ mice showing the increased number of dsred-positive interneurons (control: N = 13, MIA: N = 17, depleted: N = 13). Scale bar equal = 150 µm. **(B)** Comparison of tdTomato-positive interneurons density within all cortical layers between controls, MIA and depleted P20 mice (control: N = 13, MIA: N = 17, depleted: N = 13). **(C)** Scheme of the recording configuration and representative voltage traces of a fast-spiking PV interneuron (FS) (as in Figure 1C). **(D)** Representative average traces of inhibitory postsynaptic currents (IPSC_1-4_) evoked in PC by trains of action potentials elicited in FS interneurons in coronal cortical slices of control, MIA and microglia-depleted P20-25 mice. **(E)** Comparison of the IPSC_1_ amplitude (control: N = 13, n_cells_ = 31; MIA: N = 13, n_cells_ = 26; depleted: N = 7, n_cells_ = 17). **(F)** Comparison of the IPSC amplitude evolution during the train between the three experimental conditions. Data are represented as mean ± SEM. Two-sided unpaired Mann-Whitney test was performed to assess differences. **p < 0.01, ***p < 0.001. PC, principal cells. See also Figure S2.

### Miswiring in juveniles impairs propagation of sensory information

To examine how modifications of PV interneurons in juveniles might impact onto the functioning of cortical circuits, we first tested their role in gating incoming sensory information arriving from the thalamus. To this aim, we measured FFI in the layer 4 of the barrel cortex (**Figure S4**). Using thalamocortical slices (Agmon and Connors, 1991) that preserve the thalamic projection to the somatosensory barrel cortex, we could record in PC the monosynaptic excitatory and disynaptic inhibitory currents evoked by thalamic stimulation (**Figure S4**). These measures constitute a read out of the functional integration of PV interneurons in thalamocortical circuits (Feldmeyer et al., 2018). We found a trend for enhanced inhibitory currents triggered by thalamic stimulation in PC of both MIA and depleted pups compared to controls, whereas excitatory currents were largely unaffected (**Figure S4B**). While these experiments are relatively variable and can be biased by the angled sectioning needed to preserve thalamocortical connections, there are consistent with the enhanced inhibitory drive found in paired recordings at P20 (**Figure 2**). They also suggest, because disynaptic inhibition is more moderate than inhibition triggered by direct stimulation of interneurons (paired recordings), the existence of potential compensation mechanisms in the recruitment of PV interneurons by thalamic inputs. Nevertheless, we observed an overall increase in the Inhibition/Excitation ratio (GABA/AMPA) after MIA and prenatal depletion (**Figure S4C**), thus confirming an alteration of layer 4 PV inhibitory circuits in the context of thalamic stimulation.

Axonal arborization of most FS-PV cells is usually confined to the home-barrel, with the exception of minor populations of basket cells (Koelbl et al., 2015; Li and Huntsman, 2014; Rudy et al., 2011; Shigematsu et al., 2018). To explore whether such spatial pattern of functional projections of developing PV cells was also affected by MIA or embryonic microglial depletion, we used *PV*^*cre/+*^*;R26*^*Chr2/+*^ mice, and measured the inhibitory drive evoked by a local optogenetic activation of PV interneurons on single principal cells belonging to the same vs neighboring barrels (**Figure 3A-F**). In controls, in agreement with previous reports (Katzel et al., 2011), optogenetic stimulation of PV cells triggered robust inhibition within the same barrel albeit a very limited horizontal inhibition in adjacent barrels (**Figure 3A-F**). In both MIA and embryonically depleted juveniles, the inhibition evoked by the optogenetic stimulation of same barrel was larger than in controls (**Figure 3A** and **3D**), consistent with the larger amplitude of intra-barrel unitary synaptic responses observed in paired recordings (**Figure 2**). In addition, the inhibition onto the adjacent barrel or two barrels away was also much larger in these animals, reaching levels of responses that are obtained in controls by stimulating the barrel containing the recorded neuron (**Figure 3B-C, E-F**). Thus, in addition to a local enhanced inhibition, we detected a stronger horizontal inhibition between adjacent barrels (**Figure 3G**), revealing a miswiring of inhibitory circuits after prenatal immune challenges.

**Figure 3.**
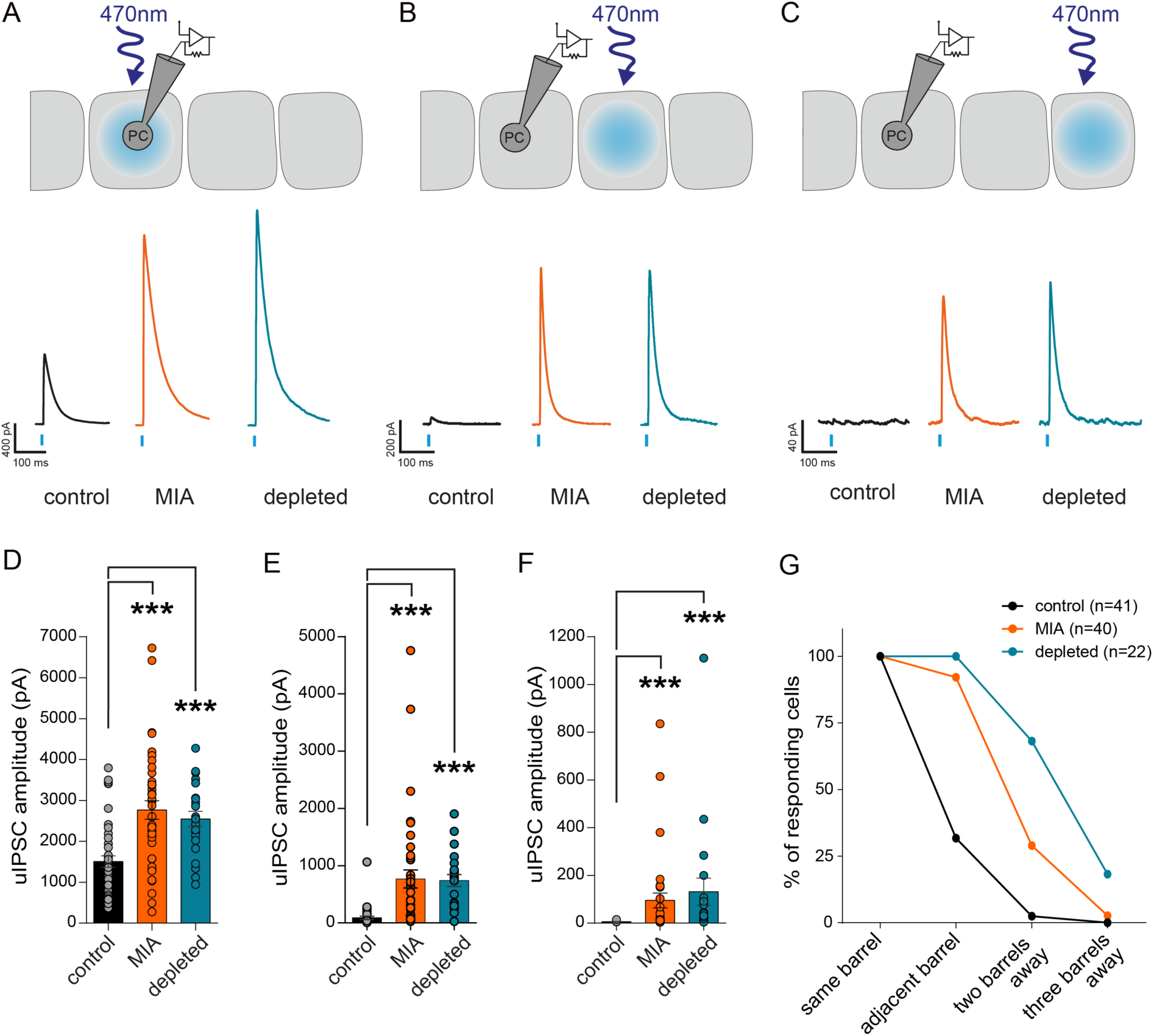
Maternal immune activation and *in utero* microglia depletion enhance inter-barrel inhibition in the cortex of juvenile mice. Representative traces of population IPSCs evoked in single principal cell (PC) by blue light stimulation of ChR2-expressing PV interneurons of the same **(A)**, of the adjacent and of the next adjacent **(C)** barrels in *PV*^*cre/+*^*;R26*^*Chr2/+*^ P20-25 mice as illustrated on the drawings. Comparison of the light-evoked IPSCs in the same **(D)**, adjacent **(E)** and next adjacent **(F)** barrels in the three experimental conditions (control: N = 18, n_cells_ = 41; MIA: N = 13, n_cells_ = 40; depleted: N = 5, n_cells_ = 22). For the next adjacent barrel, only cells displaying responses in the adjacent barrel were recorded: N = 11, n_cells_ = 16; MIA: N = 13, n_cells_ = 34; depleted: N = 5, n_cells_ = 20). **(G)** Percentage of PC responding to the optogenetic stimulation of the different barrels in control and prenatal MIA or microglia-depleted mice. Data are represented as mean ± SEM. Two-sided unpaired Mann-Whitney test was performed to assess differences. ***p < 0.001.

To further tackle whether these local circuits deficits might impair or modify the flow of sensory information *in vivo*, we visualized in real time cortical activity evoked by whisker stimulation within the barrel field of anesthetized mice by means of voltage sensitive dye imaging (**Figure 4A**) (Ferezou et al., 2006). In control conditions, as well as after MIA or transient embryonic depletion of microglia, a single whisker stimulation evoked first a signal in the corresponding column of the barrel cortex that then propagates horizontally to cover the whole barrel field (**Figure 4B** and **Movie S1**). Comparing the amplitude of the whisker-evoked signals from a region of interest matching the column corresponding to the stimulated whisker (central region of interest, ROI 1), either at the peak of the response, or at different time points after the stimulation, we observed no significant difference between mice in the control group and the MIA or the depleted group, respectively (Data not shown). However, while the latency of the response appeared similar in all groups when measured in this central ROI 1, albeit a bit shifted, MIA or transient embryonic depletion of microglia led to a significant slowdown of the lateral propagation of the signal as compared to control situation (**Figure 4B-E**). Thus, horizontal propagation of sensory information in the barrel cortex is impaired by prenatal immune challenges, consistently with enhanced horizontal inhibition across barrels.

**Figure 4.**
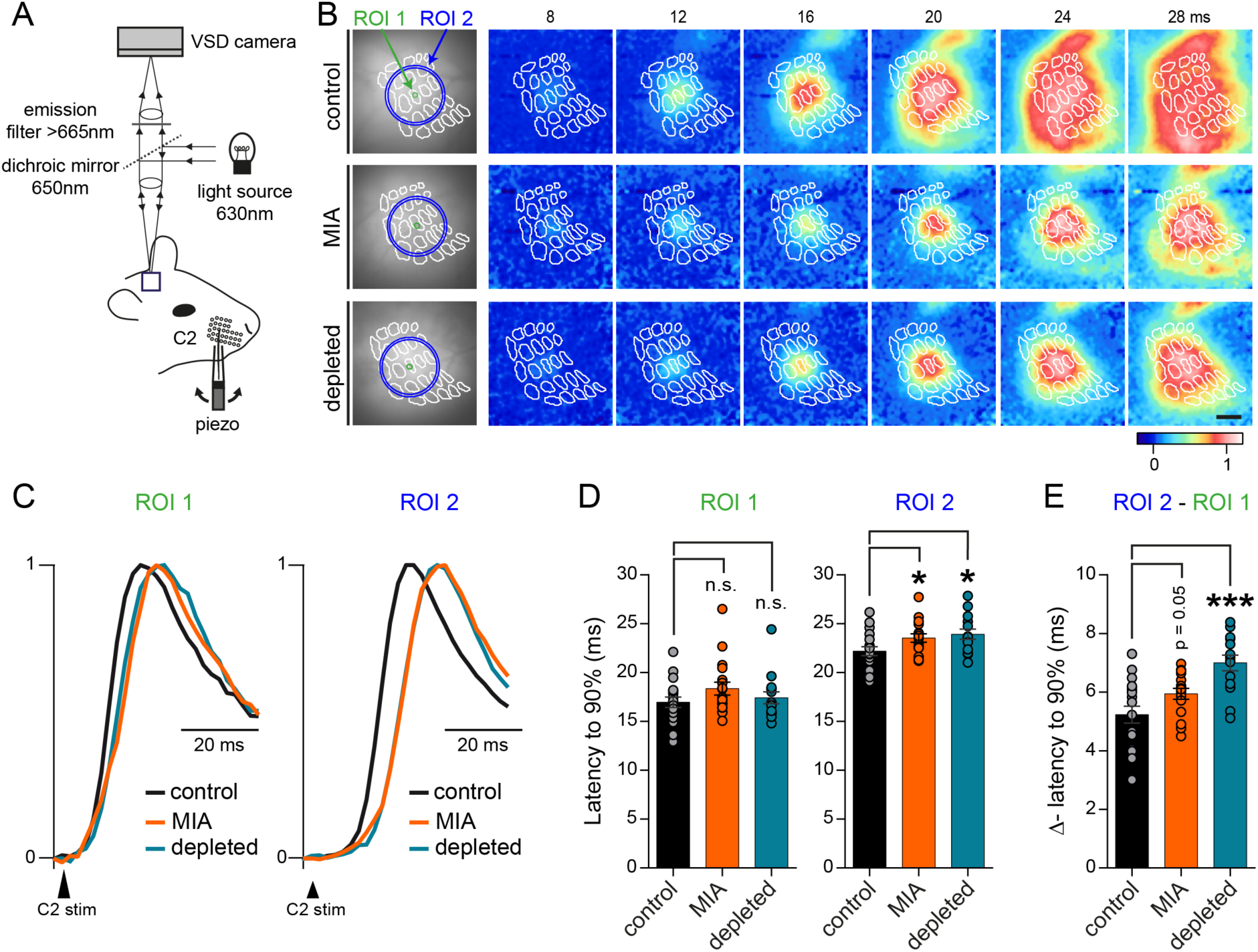
Maternal immune activation and *in utero* microglia depletion delay lateral spread of sensory information *in vivo*. **(A)** Scheme of the recording configuration. **(B)** Spatio-temporal dynamics of membrane potential changes imaged using voltage-sensitive dye (VSD) in anesthetized control, MIA and depleted animals after a brief deflection of a single whisker. The snapshots display the averaged fluorescence signal (n = 20 trials) for representative cases at six different timings (8, 12, 16, 20, 24 and 28 ms) relative to the time of deflection of the C2 whisker. The barrel map reconstructed from a *post hoc* cytochrome oxidase staining and overlaid onto the VSD signals is shown as white outlines. C2 deflection evokes a rapid localized depolarization limited to the C2 cortical barrel column and over the next milliseconds, the depolarization spreads across the barrel field. This spread of the sensory responses was assessed by quantifying the fluorescent signals from two different defined areas (region of interest (ROI) 1: green circle at the center of the column matching the stimulated whisker, and ROI 2: pixels included between the two blue circles at 550 and 600 µm radial distance from the center of ROI 1, respectively). Scale bar equal = 500 µm. **(C)** Spread of the sensory responses in ROI 1 and ROI 2 in control, MIA and depleted mice corresponding to (B). **(D)** Latencies to 90% of the peak response (averaged over 20 stimulations) computed for each ROI in control, MIA and depleted mice (control: N = 6; MIA: N = 6, depleted: N = 5; 3 whiskers stimulated by animal). **(E)** Differences of latencies between ROI 2 and ROI 1 for control, MIA and depleted mice. Data are represented as mean ± SEM. Two-sided Student’s t-test was performed to assess differences, or if the normality test failed, two-sided unpaired Mann-Whitney test was performed. *p < 0.05, ***p < 0.001, n.s., not significant. See also Movie S1.

## DISCUSSION

Our work reveals a remarkable biphasic impact of two distinct immune challenges on the normal assembly of cortical circuits, thereby providing novel insights onto how microglia dysfunction or immune risks lead to pathological brain wiring. Specifically, we report that both MIA and embryonic macrophage depletion trigger a transient increase in the number of PV neurons and of their inhibitory drive onto PC in the barrel cortex, before inducing a hypo-inhibition in the adult offspring.

The features and the temporality of the phenotypes observed in both models are remarkably similar, raising the possibility that the defect we characterized in MIA offspring results from an inability of microglia to fulfill their normal physiological roles. This is in sharp contrast with the general assumption that MIA, by perturbing microglial activity, triggers an aberrant excessive response of these immune cells (Estes and McAllister, 2016; Hirbec et al., 2018; Mattei et al., 2017; Patterson, 2011; Pont-Lezica et al., 2014). Prenatal depletion of macrophages can indeed result in changes that are opposite to those induced by MIA, such as microglial control of dopaminergic axon outgrowth, suggesting that in this context MIA does trigger an increased microglial activity (Squarzoni et al., 2014). These observations reinforce the notion that microglia exert diverse functions which are precisely regulated and determined by their environment during normal and pathological development (Casano and Peri, 2015; Mosser et al., 2017; Thion and Garel, 2017; Thion et al., 2018).

In the barrel cortex, we found that both MIA and macrophage embryonic depletion affect inhibitory circuits from early stages, thereby highlighting the importance of examining the roles of immune cells and cytokines in the development and maturation of GABAergic networks. Notably, we found that both MIA and transient embryonic macrophage depletion alter the inhibitory drive provided by PV interneurons, likely through a complex mechanism that might include changes in synapse numbers and functionalities. Our results in adult offspring are in good agreement with previous studies in other MIA models and with functional analyses in mice. In particular, several studies reported that hypo-inhibition from PV neurons constitutes a core functional deficit of neurodevelopmental disorders, impairing the normal functioning of circuits. Accordingly, modulating the activity of these cells is sufficient to improve behavioral deficits in mouse models of neurodevelopmental disorders (Canetta et al., 2016; Nelson and Valakh, 2015; Shin Yim et al., 2017). Microglia deficits, however, have been mostly associated with the dysregulation of excitatory synapse development and elimination (Hoshiko et al., 2012; Paolicelli et al., 2011; Schafer et al., 2012; Tremblay et al., 2010; Weinhard et al., 2018; Zhan et al., 2014). Our results underline that examining inhibitory synapses and their developmental trajectories will be essential to grasp the full spectrum of functional deficits associated with microglia and neuroimmune dysfunctions.

In particular, our work reveals a striking biphasic impact of prenatal immune challenges onto PV networks. In juveniles, our slice analyses and *in vivo* recordings reveal an initial excess inhibition in layer IV, an increased FFI, and a deficit in the horizontal propagation of sensory information within upper layers of the barrel cortex. These results are consistent with the existence of a strong brake on the integration of sensory inputs, at least during this period of increased functional inhibition of the PV network. The transition from juveniles to the adult situation could in theory result from an excessive, misadjusted homeostatic compensation of the large excess of inhibition we observed in juvenile mice. For instance, the excessive numbers of PV interneurons that we observe at P20 are likely eliminated by P60, since interneurons are permanently labelled by the *PV*^*cre*^ driver. In addition, the unitary IPSC are drastically reduced in MIA and depleted offspring between P20 and P60, suggesting a profound remodeling of PV interneuron connectivity. While our quantitative analysis of synaptic boutons suggests a consistent reduction in the densities of PV synapses onto the soma of PC in our experimental models between P20 and P60, these modest changes cannot account for the major electrophysiological switch that we have identified. Further experiments will be required to examine potential modifications in the synaptic strength and properties.

Irrespective of the underlying mechanisms, our results have major implications for our understanding of brain wiring. First, they highlight that inhibitory networks have several waves of adaptation to an initial embryonic perturbation, which can be drastically distinct in juveniles and adults. It thus underlines the importance of examining the full developmental trajectory of networks to understand normal physiology as well as to design adapted interventional therapies in children or adolescents. Second, our results show that prenatal immune challenges perturb the wiring of the somatosensory barrel cortex. While many studies report deficits in sensory perception associated with neurodevelopmental disorders, including ASD and SZ patients (Leekam et al., 2007; Markram and Markram, 2010), a limited number of studies focuses on the impairment associated with the circuits involved in sensory perception. Remarkably, it is well established that peripheral perturbations affecting sensory perception are sufficient to induce a large panel of behavioral deficits, including abnormal social interactions (Orefice et al., 2016). Our finding that the somatosensory cortex is drastically altered early in development, with likely consequences on sensory integration, underlines the importance of examining these structures, in addition to the better characterized alterations in the prefrontal cortex.

In conclusion, our study reveals that defects in cortical inhibition observed in adult offspring after MIA result from a complex set of events that includes an unexpected and large excess of cortical inhibition by PV cells. These observations also strongly support a role for microglia in this impaired development of cortical circuits and shed new light on how microglia and prenatal immune challenges can impact the functional wiring of the brain in neuropsychiatric disorders.

## ACKNOWLEDGMENTS

We are grateful to members of the Garel and Audinat labs for discussions and critical comments on the manuscript. We thank the IBENS Imaging Facility (France BioImaging, supported by ANR-10-INBS-04, ANR-10-LABX-54 MEMO LIFE and ANR-11-IDEX-000-02 PSL* Research University, “Investments for the future”). This work was supported by grants from INSERM, CNRS, the ERC Consolidator Grant NImO 616080 to S.G. and by grants from INSERM, the Fondation pour la Recherche Médicale (FRM: DEQ20140329488), European commission (H2020-MSCA-722053 EU-GliaPhD) to E.A.. C-A.M. was supported by the Région Île-de-France (DIM Cerveau et Pensée) and by the FRM (FDT20170739030). F.G. is an EMBO YIP awardee and is supported by Singapore Immunology Network (SIgN) core funding as well as a Singapore National Research Foundation Senior Investigatorship (NRFI) NRF2016NRF-NRFI001-02. We are grateful to C. Auger, A. Delecourt, E. Touzalin, D. Valera, C. Le Moal, A. David and M. Omnes, for excellent technical assistance.

## AUTHOR CONTRIBUTIONS

Conceptualization, M.S.T., C.A.M., S.G. and E.A.; Methodology, M.S.T., C.A.M, S.G. and E.A.; Formal analysis, M.S.T., C.A.M. and I.F.; Investigation, M.S.T., C.A.M., I.F., P.G. and S.B.; Resources, D.L., F.G., S.G. and E.A.; Writing – Original draft, M.S.T., C.A.M., I.F., S.G. and E.A.; Writing – Review & Editing, M.S.T, C.A.M., I.F., S.G. and E.A.; Visualization, M.S.T., C.A.M. and I.F.; Supervision, S.G. and E.A.; Project administration, M.S.T., S.G. and E.A.; Funding acquisition, F.G., S.G. and E.A..

## STAR METHODS

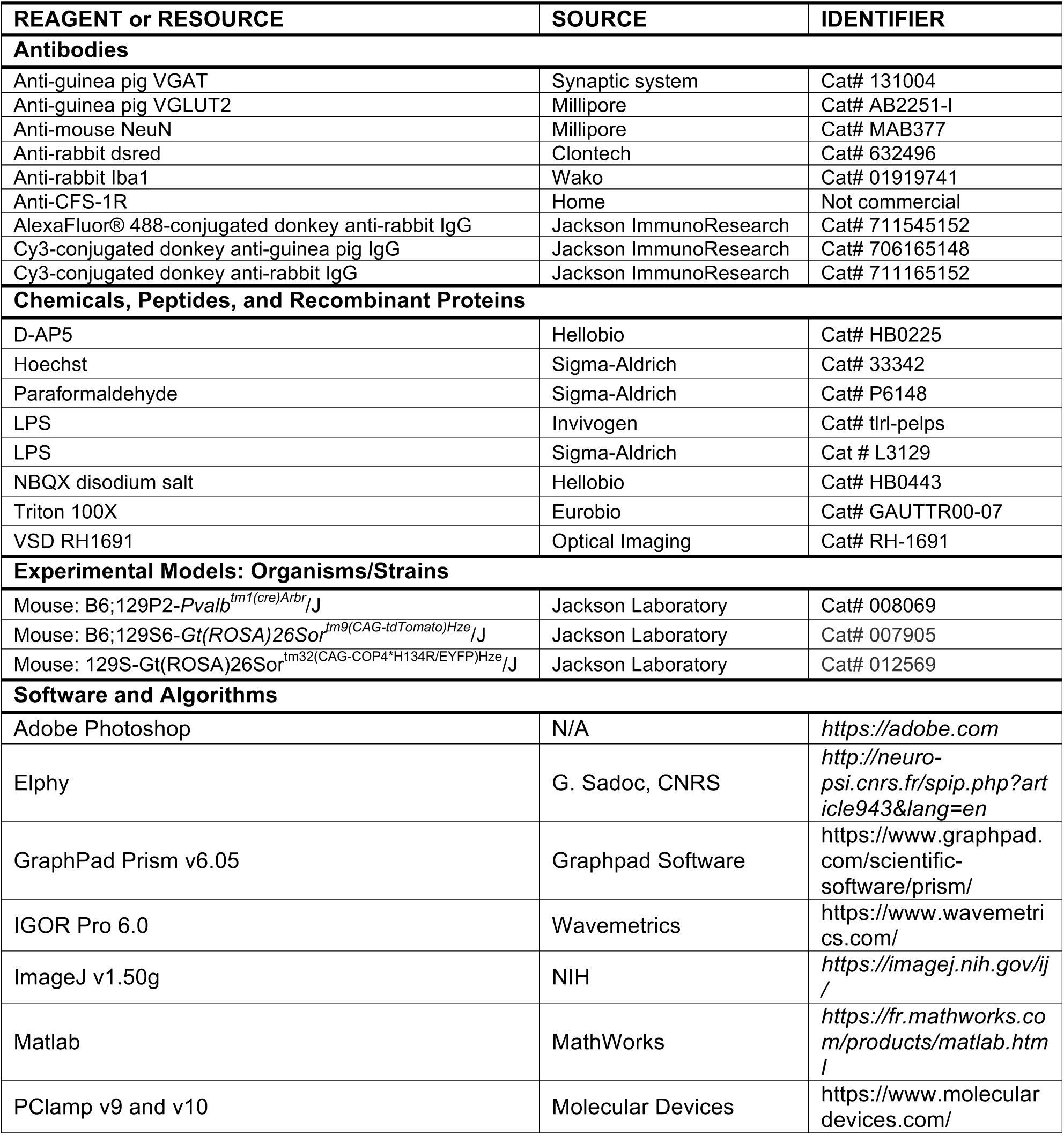
KEY RESOURCES TABLE.

### CONTACT FOR REAGENT AND RESOURCE SHARING

Further information and requests for resources and reagents should be directed to, and will fulfilled by, the Lead Contact Sonia Garel.

### EXPERIMENTAL MODEL AND SUBJECT DETAILS

#### Animals

For visualizing PV interneurons, we crossed *PV*^*Cre/+*^(Hippenmeyer et al., 2005) and *R26*^*mt/mt*^ mice (*ROSA*^*loxP-stop-loxP-tdTomato*^)(Madisen et al., 2010), resulting in the expression of tdTomato in the Parvalbumin-expressing (PV) interneurons in the offspring. For the optogenetic recordings, *PV*^*Cre/Cre*^ mice were crossed with *R26*^*Chr2/Chr2*^ floxed mice (B6; 129S-Gt(ROSA)26Sor^tm32(CAG-COP4*H134R/EYFP)Hze^/J)(Madisen et al., 2012), resulting in the expression of Channel Rhodopsin (ChR2H134R) and eYFP in PV-expressing interneurons. Both females and males have been used for all experiments presented this study, except for the voltage-sensitive dyes recording that were only performed in males. Embryonic day (E) 0.5 was set as the day of vaginal plug formation on the dam, with postnatal day (P) 0 defined as the day of birth. Mice were housed at the “Institut de Biologie de l’ENS” (IBENS), at the “Centre universitaire des Saints Pères” of Paris Descartes University, Paris, France, and at the Institut des Neurosciences de Paris Saclay” (NeuroPSI), Gif-sur-Yvette, France. These mice were handled in accordance with European regulations following the recommendations of the local ethics committees. Experiments were performed in conformity with the European (2010/63/UE) and French legislations relative to the protection of animals used for experimental and other scientific purposes (authorization numbers: 2012-0068 delivered by the local ethical committee Charles Darwin for IBENS and #59 for CEEA34.EA.027.11 and CEEA16-032 for Paris Descartes).

## METHODS DETAILS

### Microglial embryonic depletion and LPS prenatal injections

Pregnant females were administered anti-CSF-1R monoclonal antibodies (αCSF-1R, cline AFS98) intraperitoneally at E6.5 and E7.5 (3 mg per day in sterile PBS)(as in Squarzoni et al. 2014). The efficiency of the procedure was assessed by immunohistochemistry at P0. Pregnant females were administered LPS in sterile PBS (0.12 µg/g mouse; InvivoGen and Sigma) at E13.5 by a single intraperitoneal injection. Control pregnant females were injected at E13.5 with sterile PBS and showed no detectable phenotype. Three different batches of LPS were used and produced similar results.

#### Immunohistochemistry

Intracardiac perfused and dissected brains were fixed in 4% PFA at 4°C overnight. Immunohistochemistry was performed on 40 µm thick free-floating vibratome sections, as previously described (Squarzoni et al., 2014). Sections were blocked for 2 hours with PBS containing 10% fetal bovine serum and 0.01% TritonX-100, and then incubated overnight with primary antibodies. Sections were rinsed twice in PBS containing 0.01% TritonX-100, followed by several PBS washes, before overnight incubation with secondary antibodies (1/400 in PBS). Hoechst (1/1000) was used for fluorescent nuclear counterstaining.

#### Image acquisition

To determine cells density, images of the somatosensory neocortex were acquired at 10X with the Leica TCS SP5 confocal microscope. ImageJ and Adobe Photoshop were used for image processing.

### Electrophysiology

#### Slice preparation

Mice aged between 20 and 25 days (juveniles) or between 60 and 65 days (adults) were anesthetized with isoflurane, decapitated, and slices through the somatosensory cortex were cut using a Leica VT1200 vibratome in an oxygenated (5% CO2 and 95% O2) ice-cold protective extracellular solution containing for the juvenile mice (in mM): 215 sucrose, 2.5 KCl, 1.3 NaH_2_PO_4_, 25.9 NaHCO_3_, 20 D-glucose, 5 sodium pyruvate, 7 MgCl_2_, and 1 CaCl_2_ (pH 7.3, 320 mOsm), or for the adult mice (in mM): 93 NMDG, 2.5 KCl, 1.2 NaH_2_PO_4_, 30 NaHCO_3_, 20 HEPES, 2 thiourea, 25 D-glucose, 5 sodium ascorbate, 3 sodium pyruvate, 10 MgCl_2_, and 0.5 CaCl_2_ (pH 7.3, 315 mOsm). Thalamocortical slices were cut using the method described by Agmon and Connors (Agmon and Connors, 1991). After cutting, slices of young animals were incubated for 30 min at 34°C in regular artificial cerebro-spinal fluid (aCSF; pH 7.3, 310 mOsm) containing (in mM): 126 NaCl, 2.5 KCl, 26 NaHCO_3_, 1.25 NaH_2_PO_4_, 20 mM D-glucose, 1 sodium pyruvate, 1 MgCl_2_, and 2 CaCl_2_. For older animals, slices were first incubated in the same protective NMDG-containing extracellular solution for 7 min at 34°C and then incubated at 34°C for 30 min in regular aCSF. The slices were then maintained at room temperature (RT, 22-24°C) for 0.5–6h oxygenated aCSF until the recordings. After the incubation, individual slices selected for best visibility of the barrel field in cortical layer 4 were transferred to a recording chamber perfused with the same solution at 5 ml/min at RT.

#### Paired recordings

Coronal cortical slices were prepared from *PV*^*Cre/+*^*;R26*^*mt/+*^ juvenile mice, thalamocortical slices were prepared from *PV*^*Cre/+*^*;R26*^*mt/+*^ adult mice. To establish a paired recording, a whole-cell recording from a dtTomato-positive interneuron in layer 4-barrel cortex was first obtained and then sequential whole-cell recordings were made from neighboring stellate cells within 50-100 μm. The presynaptic and postsynaptic cells were patched with the above described K- and Cs-based internal solution, respectively. Presynaptic PV interneurons were held in voltage clamp at -70 mV and induced to fire single action potentials by 4 depolarizing pulses of 50 mV of 1 ms at 20 Hz. Postsynaptic cells were held in voltage-clamp at 0 mV. Paired recordings were performed in presence of the AMPA receptor antagonist (NBQX; 10 µM, Hellobio) and the NMDA receptor antagonist (D-AP5; 50µM, Hellobio).

#### Optogenetic stimulations

Coronal cortical slices were prepared from *PV*^*Cre/+*^; *R26*^*Chr2/+*^ juvenile mice. Whole-cell patch-clamp recordings were obtained from excitatory principal cells (PC) and Parvalbumin-expressing (PV) fast-spiking interneurons of the layer 4 of the somatosensory barrel cortex in voltage- and current-clamp modes, respectively. For voltage-clamp recordings patch pipettes (3-5 MΩ) were filled with a solution containing (in mM): 125 CsMeSO_3_, 10 HEPES, 10 EGTA, 8 TEA-Cl, 5 4-AP, 0.4 GTP-Na, 4 ATP-Na_2_, 1 CaCl_2_ and 1 MgCl_2_ (pH 7.3, 290 mOsm). For current-clamp recordings aiming at adjusting parameters of the optogenetic stimulation (see below), pipettes (3-5 MΩ) were filled with a solution containing in mM: 125 K-Gluconate, 2 MgCl_2_, 10 HEPES, 0.4 Na^+^-GTP, 4 ATP-Na_2_, 10 Phosphocreatine disodium salt, 5 KCl, 0.5 EGTA (pH 7.3, 290 mOsm). Recordings were performed in the presence of the following glutamatergic receptors blockers: 10 µM 2,3-dioxo-6-nitro-1,2,3,4-tetrahydrobenzo[f]quinoxaline-7-sulfonamide disodium salt (NBQX; Hellobio) and 50 μM D-(–)-2-amino-5-phosphonopentanoic acid (D-AP5; Hellobio). IPSCs recorded in layer 4 PC were optogenetically evoked at 0 mV by driving the ChR2 activity in PV interneurons with 470 nm blue light (0.9 mW/mm_2_) delivered by a LED (Cairn Research, OptoLED) through the 40X water-immersion objective (Olympus). We used a 100 µm diameter light field (*i*.*e*. about the size of one barrel) and light stimulations of 5 ms were delivered at 0.05 Hz. In pilot experiments, the light intensity was adjusted to 1.5 times the threshold for eliciting one action potential in FS cells and we verified that this threshold did not differ among the different experimental groups (data not shown).

#### Thalamic stimulations

Thalamocortical EPSCs and IPSCs were evoked in principal neurons of the barrels using a stainless steel bipolar microelectrode (100 µm tip diameter, 250 µm intertip distance; Rhodes Medical Instruments) connected to a stimulus isSolation unit (Iso-stim 01D; npi Electronic) and placed in the ventrobasal thalamus or in the internal capsule near the thalamus border. The intensity of stimulation (0.1 ms at 0.05 Hz) was first adjusted according to the protocol of minimal stimulation described in (Hoshiko et al., 2012) and then increased 1.5 times to ensure the recruitment of feedforward inhibition. The EPSCs and IPSCs were recorded at -70 mV and 0 mV, respectively.

### Voltage-sensitive dye (VSD) imaging

VSD imaging of the cortical activity evoked by tactile whisker stimulation was performed on P27-P33 male mice under isoflurane anesthesia (induction: 3-4%, maintenance: 1-1.5%). The left barrel cortex was exposed and stained for 1h with the VSD RH1691 (Optical Imaging Ldt, Israel) at 1mg/ml, in Ringer’s solution containing [in mM]: 135 NaCl, 5 KCl, 5 HEPES, 1.8 CaCl_2_, 1 MgCl_2_. After removing the unbound dye, the cortex was covered with agarose (0.5-1%) and a coverslip. Cortical imaging was performed through a tandem-lens fluorescence microscope (SciMedia), equipped with one Leica PlanApo 5x (objective side) and one Leica PlanApo 1x (condensing side), a 100 W halogen lamp gated with an electronic shutter, a 630 nm excitation filter, a 650 nm dichroic mirror, and a long-pass 665 nm emission filter (**Figure 4A**). The field of view was 2.5 x 2.5 mm, resulting in a pixel resolution of 25 x 25 µm.

The right C2, C3 or C4 macrovibrissae were individually deflected in a pseudo-randomized manner using a multi-whisker stimulator (Jacob et al., 2010). Whiskers were inserted keeping their natural angle in 27G stainless steel tubes attached to the piezoelectric benders (Noliac, Denmark), leaving 2 mm between the tip of the tube and the whisker base. Each whisker deflection consisted of a 95 µm displacement (measured at the tip of the tube) a 2 ms rising time, a 2 ms plateau and a 2 ms fall. Specific filters were applied to the voltage commands to prevent mechanical ringing of the stimulators. Following the experiments mice were perfused with saline followed by paraformaldehyde (4% in 0.1M phosphate buffer). After an overnight post-fixation in paraformaldehyde, the brains were cut in 100 µm-thick tangential sections that were stained for cytochrome oxidase. Microphotographs of the tangential sections were registered and the barrel maps reconstructed using a method implemented in Matlab (MathWorks, USA), as previously described (Perronnet et al., 2016). The functional VSD data were aligned with the reconstructed barrel maps by using the superficial blood vessels as anatomical landmarks.

### QUANTIFICATION AND STATISTICAL ANALYSIS

#### Image analysis

For barrel characterization, images were converted to greys and inverted before delineating vglut2-positive barrels (thalamocortical) and counting Iba1-positive cells. Hoechst and vglut2 stainings were used to delineate the thickness of cortical layers. The relative fluorescence intensity across the C2 barrel was assessed as described in (Toda et al., 2013). Dsred-positive cells quantification in *PV*^*cre/+*^*;R26*^*tom/+*^ mice was performed using Bitplane Imaris software. Synapses were automatically detected with the ImageJ software by first applying despeckle followed by a smooth filter, then local maxima were found on the VGAT channel. The local maximas were than visualised on the NeuN channel, and the number of maximas par soma were quantified. Results are represented by the number of synapses par soma.

#### Electrophysiological data acquisition and analysis

Whole-cell patch-clamp recordings were obtained using either Axopatch 200B or 700B amplifiers (Molecular Devices). Currents and potentials were low-pass filtered at 5 kHz or 6 kHz, collected at a 10 kHz frequency, and analyzed off-line using pClamp 10.7 (Molecular Devices) for voltage-clamp recordings, or using a custom written procedure in IGOR Pro 6.0 (Wavemetrics) for current-clamp recordings. For all recordings performed with K-gluconate or Cs-gluconate in the pipette, potentials were corrected for a junction potential of -10 mV. Series resistances (Rs) were not compensated but continuously monitored using 5 or 10 mV hyperpolarizing pulses. Recordings were discarded if Rs increased by >20% during the whole experiment. For all experiments, the amplitude of EPSCs and IPSCs were quantified from the average of 15–30 consecutives responses. For paired-recording analysis, the peak responses to each stimulation were measured and then averaged over all trials. The paired-pulse ratio was calculated as the average peak amplitude of the second response divided by that of the first one. In current clamp experiments aiming at characterizing basic membrane properties, a series of hyperpolarizing and depolarizing current steps were applied for 1 s in 25-50 pA increments at 5 s intervals. Pairs obtained with presynaptic cells displaying low firing rate or adaptation ratio (<0.6) or small AHP (<10 mV) were discarded.

#### Voltage-sensitive dye data acquisition and analysis

Using a CMOS-based camera (MiCam Ultima, SciMedia), sequences of images of 1 s were acquired at 500 Hz, with or without (blank trials) whisker stimulation, alternately, every 20 s. Variations of the light intensity were initially recorded as variations over the resting light intensity (first acquired frame). Acquisition and data preprocessing were done using in-house software (Elphy, G. Sadoc, UNIC-CNRS), further analyses were made using custom written routines in IgorPro (WaveMetrics, USA). Subtraction of a pixel by pixel best-fit double-exponential from the averaged unstimulated sequence was used to correct for photobleaching. For each averaged whisker evoked response (n = 20 trials), a 2D-gaussian fit of the evoked signal at 12 ms following the start of the whisker stimulus was used to localize the center of a first region of interest (ROI) defined as a 100 µm-diameter circle. A second, more distal ROI, was defined as a 50 µm-wide ring placed at 575 µm from the center of the first ROI. Profiles of fluorescence were computed from these ROIs and normalized to determine the latency to 90% of the peak. Responses with amplitudes < 0.1% were discarded from the analysis. Signals measured in response to the deflection of the 3 individual whiskers were considered as independent.

#### Statistics

All data are presented as mean ± SEM. Non-parametric two-sided Mann-Whitney *U*-tests were used to compare two distributions. If the normality test was statistically significant, two-sided Student’s t-tests were used to compare two distributions. Two-way ANOVA with Sidak post hoc test were used to compare groups of data. All graphs and statistical analyses were generated using GraphPad Prism software. *p < 0.05, **p < 0.01, ***p < 0.001.

### DATA AND SOFTWARE AVAILABILITY

The data supporting the findings of this study are available upon request to the corresponding authors.

