## Supplemental Material for "Biphasic impact of prenatal inflammation and macrophage depletion on the wiring of neocortical inhibitory circuits"

**Thion*, Mosser* et al.**

### SUPPLEMENTAL INFORMATION

Supplemental Information includes 4 supplementary figures and 1 supplementary movie.

### SUPPLEMENTAL FIGURES

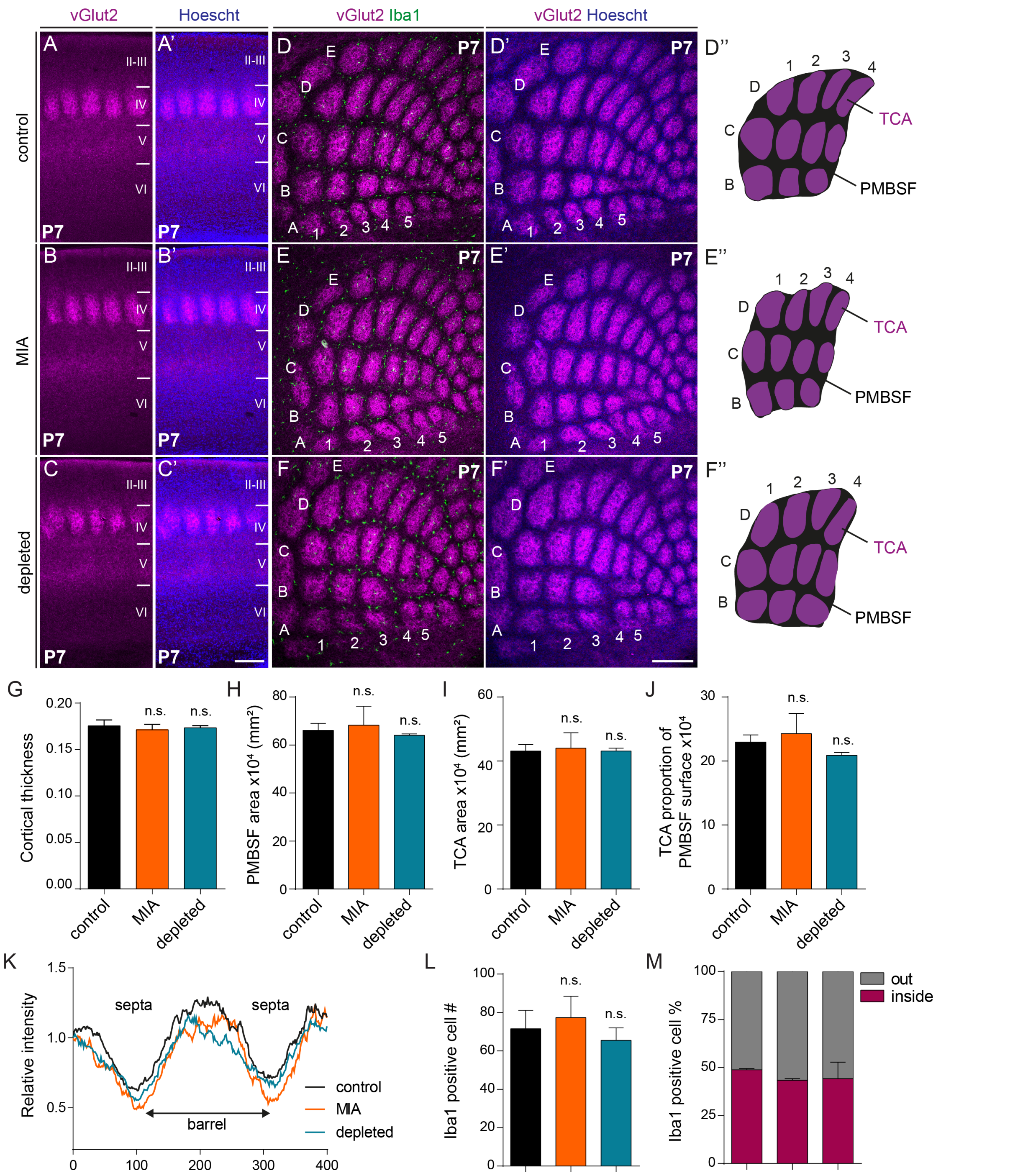


**Supplementary Figure 1.** ***In utero*** **microglia depletion and Maternal Immune Activation do not disrupt the formation of barrels, Related to Figures 1-4.**

**(A-C')** Coronal sections through the P7 posterior medial barrel subfield (PMBSF) of the primary somatosensory cortex of control, MIA and microglia depleted mice stained with vGlut2 antibodies to label thalamocortical axons (TCA) and counterstained with Hoechst. Scale bar = 200 µm. **(D-F')** Tangential sections through the P7 barrel field of control, MIA and depleted mice immunostained with vGlut2, the microglia marker Iba1 and Hoechst. Scale bar = 200 µm. **(D''-F'')** Reconstructions of the PMBSF in control, MIA and depleted P7 mice and showing no significant changes among the three conditions. **(G-J)** Quantifications of the cortical thickness **(G)**, the PMBSF area **(H)**, the TCA area **(I)** and the proportion of TCA overs PMBSF surfaces **(J)** in control, MIA and depleted P7 mice (control: N = 5, MIA: N = 3, depleted: N = 4). **(K)** Relative fluorescence intensity across the C2 barrel in control, MIA and depleted mice (control: N = 5, MIA: N = 3, depleted: N = 4). **(L-M)** Number of microglial cells within the barrel field **(L)** and the proportion of these cells located inside or outside the barrels **(M)** among control, MIA and depleted P7 mice (control: N = 4, MIA: N = 3, depleted: N = 2). Data are represented as mean ± SEM. Two-sided unpaired Mann-Whitney test was performed to assess differences. n.s., not significant.

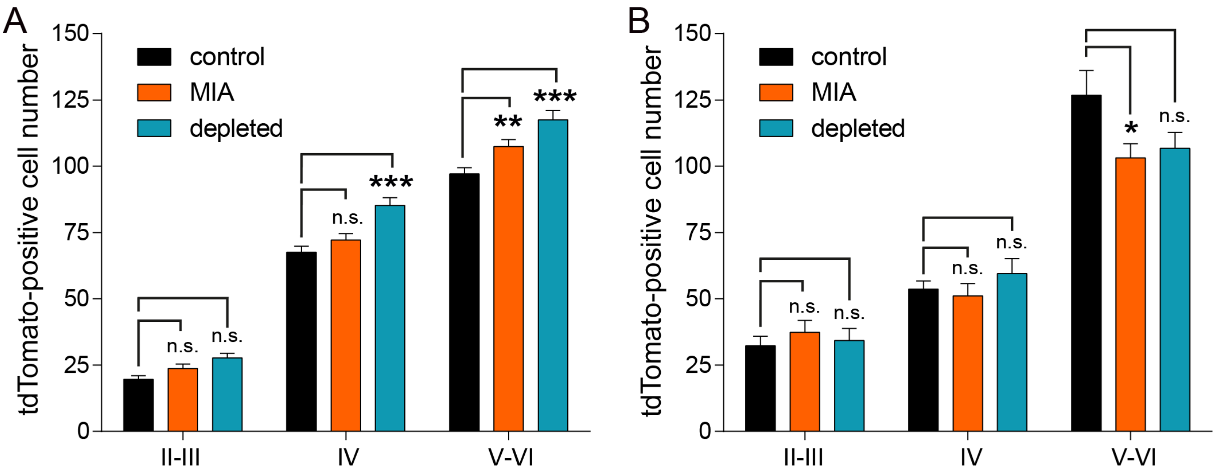


**Supplementary Figure 2. *In utero* microglia depletion and Maternal Immune Activation impair the laminar density of PV interneurons, Related to Figures 1 and 2.**

**(A)** Comparison of PV interneurons numbers within cortical upper (II-III), IV and deep (V-VI) layers between controls, MIA and depleted P20 mice (control: N = 10, MIA: N = 6, depleted: N = 4) and **(B)** P60 mice (control: N = 13, MIA: N = 17, depleted: N = 13). Data are represented as means ± SEM; two-way ANOVA with Sidak post hoc test was performed to assess differences at each stage. *$p < 0.05$, **$p < 0.01$, ***$p < 0.001$, n.s., not significant.

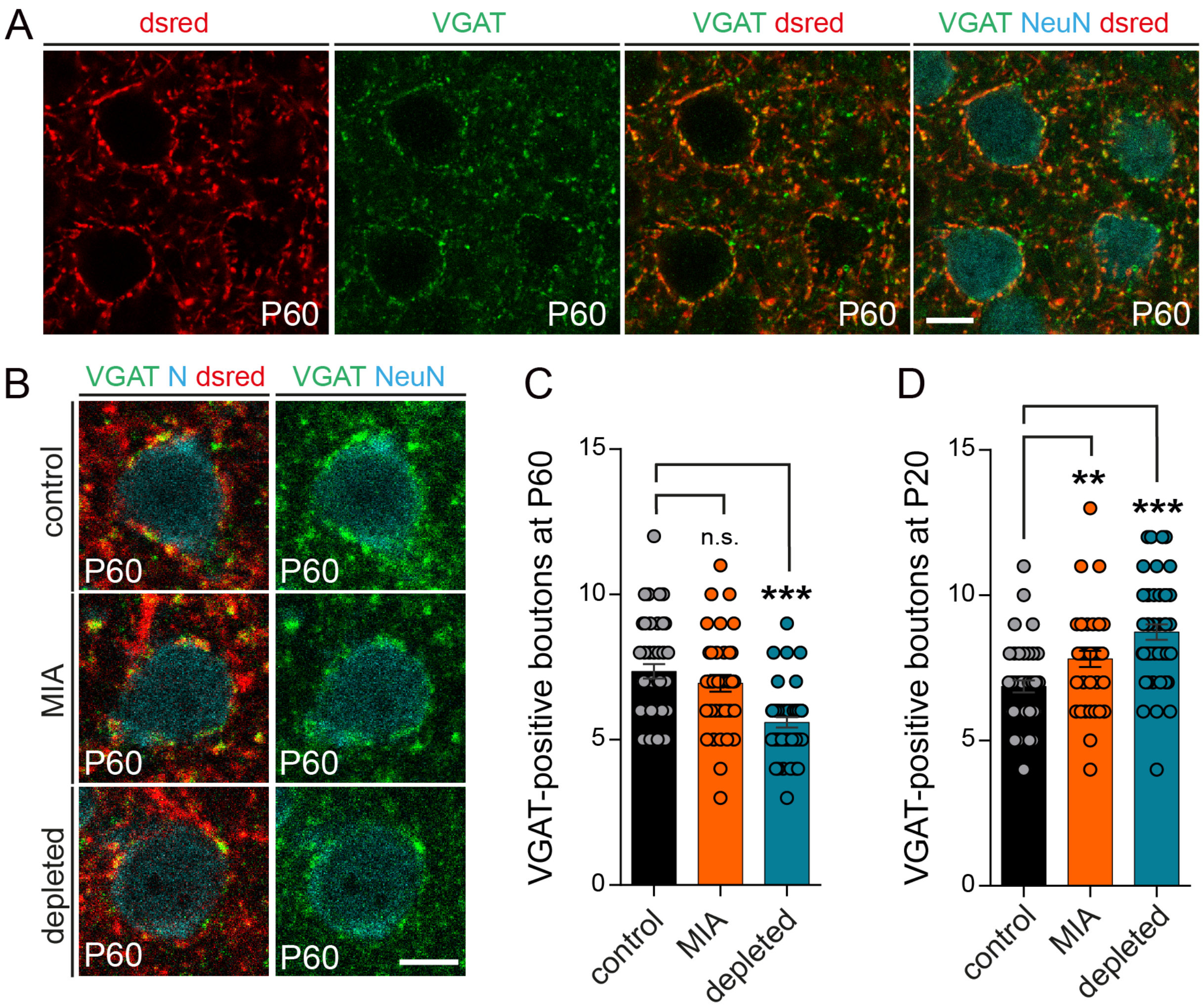


**Supplementary Figure 3. *In utero* microglia depletion and Maternal Immune Activation mildly affect the densities of synaptic boutons formed by PV interneurons onto the soma of PC, Related to Figures 1 and 2.**

**(A)** P60 primary somatosensory cortex of control *PV*$^{Cre/+}$;*R26*$^{tom/+}$ mice stained with dsred, VGAT and NeuN antibodies to respectively label PV interneurons, inhibitory synpases and neuronal nucleus and counterstained with Hoechst.

Scale bar = 10 µm. **(B)** P60 primary somatosensory cortex of primary somatosensory cortex of control, MIA and microglia depleted *PV*$^{Cre/+}$;*R26*$^{tom/+}$ mice stained with dsred, VGAT and NeuN (N) antibodies to respectively label PV interneurons, inhibitory synapses and neuronal nucleus and counterstained with Hoechst. Scale bar = 5 µm.

**(C)** Number of synaptic boutons formed by PV interneurons onto the soma of PC in the somatosensory cortex of controls, MIA and depleted P60 $PV^{Cre/+};R26^{tom/+}$ mice (N = 3 mice per conditions, at least 10 cells were analyzed per mouse). **(D)** Number of synaptic boutons formed by PV interneurons onto the soma of PC in the somatosensory cortex of controls, MIA and depleted P20 $PV^{Cre/+};R26^{tom/+}$ mice (N = 3 mice per conditions, at least 10 cells were analyzed per mouse).

Data are represented as mean ± SEM. Two-sided unpaired Mann-Whitney test was performed to assess differences. **p < 0.01, ***p < 0.001, n.s., not significant.

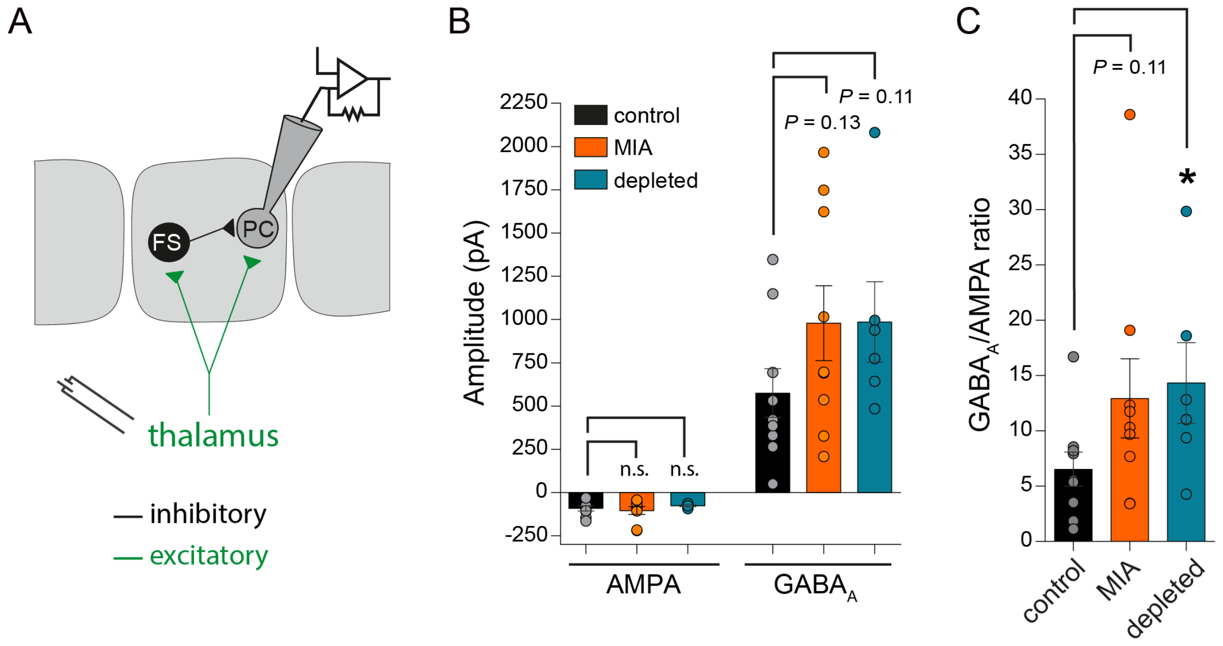


**Figure S4. Effect of Maternal Immune Activation and *in utero* microglia depletion on feed-forward inhibition in layer 4 of the juvenile barrel cortex, Related to Figure 3.**

**(A)** Scheme of the recording configuration in thalamocortical slices illustrating the stimulation electrode in the thalamus, the whole-cell recording pipette onto a layer 4 principal cell (PC) and the di-synaptic circuit recruiting fast-spiking PV interneurons in the barrel cortex. **(B)** Average amplitude distribution of the monosynaptic EPSCs (AMPA) and disynaptic IPSCs ($GABA_A$) evoked in PCs by thalamic stimulation for 8 control, 8 MIA and 6 microglia depleted mice (control: $n_{cells}$ = 23; MIA: $n_{cells}$ = 25; depleted: $n_{cells}$ = 21). EPSCs and IPSCs were recorded at – 70 mV and 0 mV, respectively. **(C)** Distribution of the average I/E ratio (or $GABA_A$/AMPA ratio) for the same mice as in (B).

Data are represented as mean ± SEM. Two-sided Student's t-test was performed to assess differences, or if the normality test failed, two-sided unpaired Mann-Whitney test was performed. *$p < 0.05$, n.s., not significant.

### SUPPLEMENTAL MOVIE

**Supplementary Movie 1. Maternal Immune Activation and prenatal microglia depletion delay the lateral spread of sensory information *in vivo,* Related to Figure 4.**
